# Tradeoffs in ATP metabolisms via hypoxic gradient migration assays

**DOI:** 10.1101/2024.04.06.588411

**Authors:** Mohamad Orabi, Kai Duan, Mengyang Zhou, Joe F. Lo

## Abstract

Migration and scratch assays are helpful tools to investigate wound healing and tissue regeneration processes, especially under disease conditions such as diabetes. However, traditional migration (injury-free) assays and scratch (with injury) assays are limited in their control over cellular environments and provide only simplified read-outs of their results. On the other hand, microfluidic-based cell assays offer a distinct advantage in their integration and scalability for multiple modalities and concentrations in a single device. Additionally, *in situ* stimulation and detection helps to avoid variabilities between individual bioassays. To realize an enhanced, smarter migration assay, we leveraged our multilayered oxygen gradient (1-16%) to study HaCaT migrations in diabetic conditions with spatial and metabolic read-outs. An analysis of the spatial migration over time observed a new dynamic between hypoxia (at 4.16-9.14% O2) and hyperglycemia. Furthermore, *in situ* adenosine triphosphate (ATP) and reactive oxygen species (ROS) responses suggest that this dynamic represents a switch between stationary versus motile modes of metabolism. Thus, elevated glucose and hypoxia are synergistic triggers of this switch under disease conditions. These findings illustrate the benefits of spatial microfluidics for modeling complex diseases such as hypoxia and diabetes, where multimodal measurements provide a more deterministic view of the underlying processes.

## 1. Introduction

### 1.1. Diabetes, Hypoxia, and Wound Healing

Diabetes and hypoxia are two factors that can hinder the wound healing process, especially when presented together. Diabetes is characterized by systemic hyperglycemia due to either insufficient insulin or insulin resistance (**Egan and Dinneen, 2019**). On the other hand, hypoxia occurs when the tissues do not receive sufficient oxygen due to decreased metabolism, hypoxemia, or poor microvasculature. Tissue hypoxia is a common complication in obesity and diabetes and impairs cell proliferation, angiogenesis, and thus wound healing (**Singh et al., 2017**). As of 2019, about 463 million adults were living with diabetes, with projections of 700 million by 2045. Given the prevalence of ulcers, chronic wounds, and amputations in the diabetic population, this represents a severe public health concern worldwide (**Jin et al., 2020)**. Both diabetes and hypoxia affect multiple stages of wound healing from inflammation to remodeling and modulate complex biological and molecular processes (**Yang et al., 2021)**. Thus, understanding the interaction between diabetes and hypoxia in the context of impaired healing is a crucial public health issue that requires immediate attention.

Wound healing involves a complex interplay of cell types including keratinocytes, fibroblasts, endothelial cells, and inflammatory cells (**Rodrigues et al. 2019; Orabi and Ghosh, 2023**). Platelets and injured keratinocytes in early wound healing release growth factors that attract inflammatory cells to the area (**Yang et al., 2021**). For example, macrophage infiltration is necessary for the cleanup and preparation of the wound bed (**Tang et al., 2021**), but their reparative functions and ability to stimulate re-epithelialization may be impaired under hyperglycemia (**Deng et al., 2021; Ezhilarasu et al., 2020**). While previous research has focused on inflammatory and other wound cells (**Wulff et al., 2012**), diabetic and hypoxic effects on keratinocytes—cells central to re-epithelialization—have not been thoroughly investigated. To address these challenges, researchers have developed microfluidic devices that allow either temporal or spatial analysis of molecular transport in HaCaT cells (**Maurya et al., 2022**). However, these devices must choose between maintaining gradients or enabling time dynamics due to limited diffusion within microchannels (**Duan et al., 2023a**). In previous work, we address the limitation on gas diffusion by leveraging nitrogen convection to drive oxygen transport, resulting in dynamic, spatial oxygen gradients. In this study, we further applied our oxygen gradient to keratinocytes under diabetic conditions, providing simultaneous cell migration, mitochondrial and cytoplasmic ATP, and oxidative stress detections at various levels of hypoxia. In doing so, we uncovered a hidden relationship between HaCaT cell migration versus oxygen modulation of mitochondrial and cytoplasmic ATP levels.

### 1.2. Modeling Diabetic Wound Healing

The HaCaT cell line is skin keratinocytes that provide barrier function for the body. The cell line can further differentiate into various skin cell types to provide models for *in vitro* dermatological and wound healing studies (**Leung et al., 2017; Colombo et al., 2017**). Re-epithelialization is a crucial step in wound healing that involves keratinocyte migration and proliferation to close the wound bed and restore epidermal barrier functions (**Figure 1a**) (**Tang et al., 2021**). HaCaT cells can be used to model different aspects of wound healing, including the effects of growth factors, cytokines, extracellular matrix components, and potential factors such as hypoxia or hyperglycemia (**Yang et al., 2020; Lee et al., 2019**). After cell injury, HaCaT releases ATP, UTP, and derivatives into the wound site (**Laberge et al., 2018**). Subsequently, ATP and UTP propagate cell signals via P2 receptors that have broad functions in cell proliferation, differentiation, cytokine signaling, and immune cell recruitment, as observed in corneal epithelial cell injury models (**McEwan et al., 2021**). On the other hand, hyperglycemia-induced reactive oxygen species (ROS) can alter signaling pathways that regulate cell migration, triggering NF-kB-based inflammation and cell adhesion, thus hindering cell migration. Tissue hypoxia also modulates the same NF-kB/COX-2 pathway, which is triggered by reactive oxygen species (ROS) (**Yang et al., 2011**).

**Figure 1.**
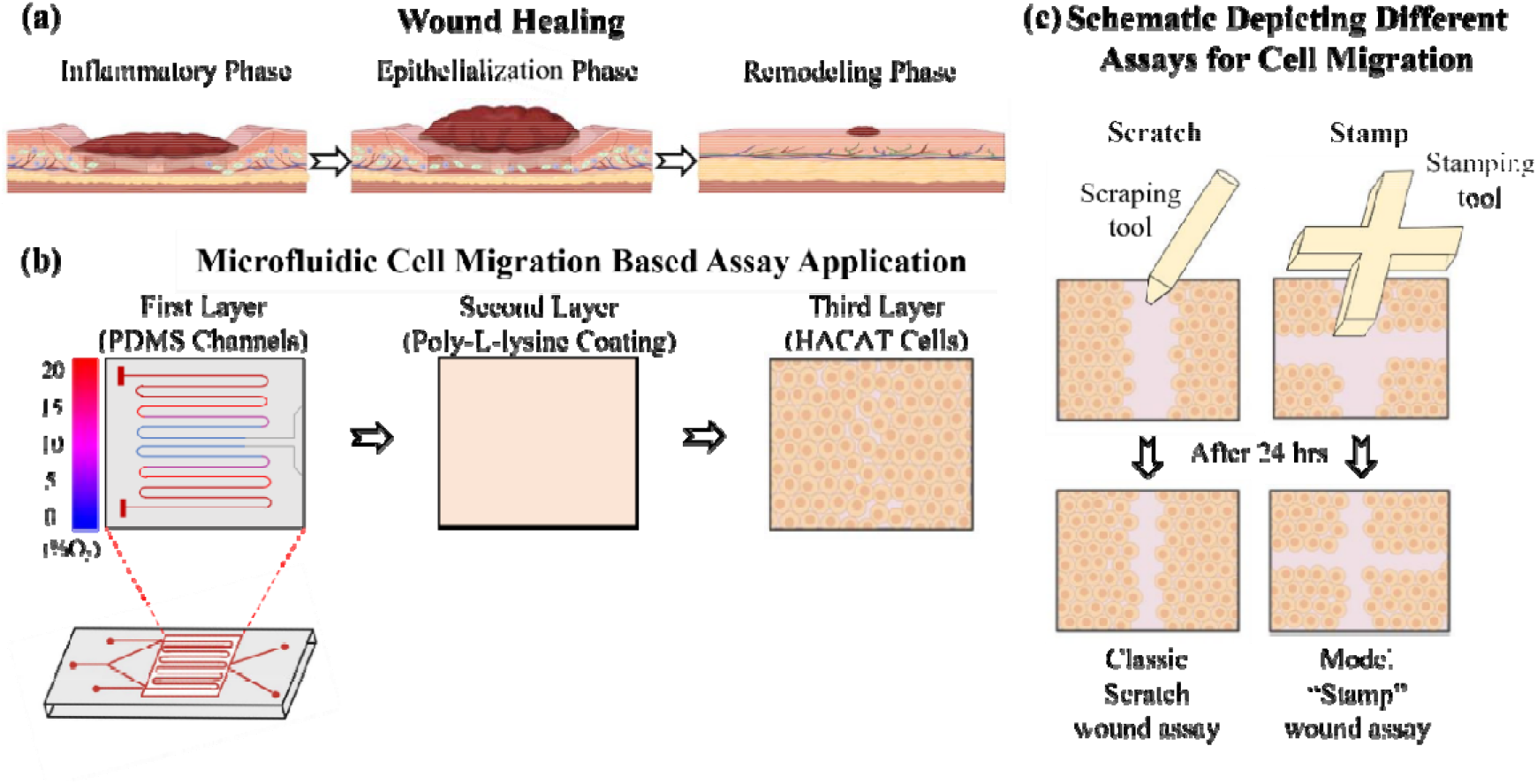
Microfluidic oxygen gradient with a spatial poly-l-lysine sensor. (a) The stages of healing are inflammation, epithelialization, and remodeling. The inflammatory phase is marked by the white blood cells, growth factors, nutrients, and enzymes that create swelling, heat, pain, and redness. For the epithelialization phase, the body begins to create a scab, which provides temporary protection for the wound. Lastly, for the remodeling phase, the body fills in the wound with new tissue to restore the skin’s integrity. (b) The microfluidic device has multiple layers. The gas layer maintains a gradient of oxygen (red) and nitrogen (blue) using a 200 μm polydimethylsiloxane (PDMS) membrane, which is permeable to oxygen and diffuses the gradient to poly-l-lysine and cell culture seeded on top. The oxygen gradient is designed to be symmetrical, so either half can be used. The PDMS layer is coated with ply-l-lysine to improve cell attachment and adhesivity. The open-top aqueous reservoir facilitates normal cell culture processes and fluorescence microscopy. (c) Moreover, cell migration of HaCaT cells was due to stamp and scratch assays. A 200 μl sanitized micropipette was used to create the scratch wound and a 3D printed stamping tool was used to create a stamp wound.

Furthermore, ROS are essential regulators of cell migration, but excessive ROS can lead to overactive Rho GTPases, impairing cell migration. Glucose-induced ROS can also damage proteins and lipids in the cell membrane and cytoskeleton, which are essential for cell movements. On the other hand, Yang et al. studied HaCaT injuries and inflammation related to the hypoxic modulation of the NF-kB/COX-2 pathway, which is also triggered by reactive oxygen species (ROS) (**Yang et al., 2011**). This illustrates the synergy between hypoxia and hyperglycemia. McEwan et al. explored the autocrine regulation of wound healing via ATP using an *in vitro* scratch wound model (**McEwan et al., 2021**). They found that ATP activates P2Y2 receptors and stimulated PLC/IP3 signaling, leading to an increase in intercellular Ca^2+^ that enables wound closure independent of cell proliferation. Kim’s and Li’s group investigated the effects of platelet-rich plasma (PRP) in treating diabetic foot ulcers (DFUs) and showed PRP regulation of NF-kB through PDCD4 plays an important anti-inflammatory role in DFUs **(Kim et al., 2013; Li et al., 2019a)**. Thus, we propose a possible synergy between hypoxia and hyperglycemia in wound cell migration through inflammation. These findings have the potential to lead to the development of new therapeutic targets for difficult-to-heal wounds.

### 1.3. Conventional Migration Assays

The gold standard in migration assay involves creating a scratch on a petri dish using a sterile pipette (**Michalczyk et al., 2018**), which disrupts the cell monolayer and generates an empty space devoid of cells (**Mouritzen and Jenssen, 2018**). Additionally, gold nanoparticle (GNP) coating aids in the quantification of cell movements, including nanoparticles functionalized with growth factors (**Leu et al., 2012)**. Functionalized GNPs have also been successfully applied to full-thickness models to promote wound re-epithelialization **(Pan et al., 2018; Li et al., 2019b)**. Alternatively, researchers have also applied migration assays without injury as a comparison. Injury-free assays such as cell stamping, Figure 1c, provide a baseline cell migration that may be affected by factors outside of wounding (**Yin et al., 2022**), for example, exposure to diabetic conditions. Sano et al. conducted a scientific investigation to examine the impact of fluctuating oxygen environments on the progression of wound healing in a murine model (**Sano et al., 2012**). An ample supply of oxygen has been reported to facilitate proper epithelialization and the formation of granulation tissue, while also promoting increased neovascularization in hypoxic conditions, suggesting a compensatory response to the hypoxic state. In contrast, microfluidic injury assays using scratching or trypsin flow focusing provide the actual wound stimulus for modeling the wounds or ulcers we are investigating in diabetes (**Lin et al., 2019; Yontem et al., 2019**). To test our hypothesis of a hyperglycemia-hypoxia synergy in diabetic wounds, we applied our oxygen gradient to glucose-exposed keratinocytes in both injury-based (scratch) and injury-free (stamp) migration assays.

### 1.4. The *in situ* Multimodal Microfluidic Migration Assay

A gradient oxygen microfluidic device was designed by combining gas microfluidics with a gas-permeable poly-L-lysine-functionalized membrane. This mechanism allowed for the manipulation of oxygen levels across a large monolayer of cells for migration assay. The gas microfluidics balanced the oxygen diffusion against nitrogen convection and created a 0-16% hypoxic gradient delivered across a 200 μm polydimethylsiloxane (PDMS) membrane (Figure 1b) (**Li et al., 2015**). The gradient was symmetrically designed so that both halves of the device could be used (**Duan et al., 2023a**). The aqueous side of the membrane enables cell exposure to glucose stimulation and *in situ* detections of ATP and ROS. The device was fabricated by bonding SU8 molded membranes, including the gas channels, layer by layer with a chamber gasket on top for cell culturing. The device offered a unique way to visualize oxygen concentrations and cell migration in HaCaT cell cultures that was not possible with contemporary migration assays. We applied both stamp and scratch assays to study HaCaT migration (Figure 1c). A 3D printed insert defined the size and shape of cell patterning in the microfluidic chamber for the stamp-based assay (**Duan et al., 2023b**). In contrast, the scratch assays applied a sterile pipette tip to manually remove a line of cells, generating a cell-free gap bordered by injured cells. The resultant device offered a unique, multimodal way to visualize oxygen concentrations and cell migration in HaCaT cultures that was not possible with contemporary migration assays.

## 2. Material and Methods

### 2.1. Microfluidic fabrication

The SU-8 photolithography technique was used to create the molds for the device. Firstly, a layer of SU-8 2100 (Microchem, USA) was applied to a silicon wafer via spin coating. This resulted in a 100 μm thick layer which was then soft baked and exposed to UV light using a mask. Following the post-exposure bake, the layer was developed with an SU-8 developer. The thickness of the gradient layer channel was determined using a surface profilometer. The wafer’s surface was treated with plasma and then left in a vacuum overnight for salinization with trichloro (1H,1H,2H,2H-perfluorooctyl).

To coat the gas microfluidics with poly-l-lysine, a PDMS membrane was created using the previously fabricated SU-8 mold. The membrane was placed on a heated plate at 95°C for 30 minutes. Following the same process, a second layer was added on top of the first layer. The resulting 200 μm membrane that was molded over the gas microchannel master was cut and then bonded to a glass slide, thereby enclosing the gas channels with the exposed textures. Lastly, a PDMS gasket, which was 1 cm thick, was cut and bonded to create the cell culture chamber.

### 2.2. Cell culture

HaCaT cells (ATCC) were cultured in regular Dulbecco’s modified Eagle’s medium (DMEM) supplemented with 10% (v/v) fetal bovine serum (FBS) and 1% (v/v) penicillin-streptomycin. Cells were incubated until confluence at 37°C and 5% CO_2_. In this study, cells up to passage 7 were used. To culture inside the microfluidics, 1 × 10^6^ HaCaT cells were loaded onto the device with 2 mL of medium without glucose. The device was incubated for 5 hours prior to the start of experiments to allow HaCaT cells to acclimatize to normal glucose levels before stimulation. Following the incubation, the media and the stamp were removed for stamp assay and a scratch was applied for scratch assay. Then they were washed 3 times with Dulbecco’s Phosphate Buffered Saline (DPBS). During the microfluidic experiments, the media was replaced with a low glucose medium (Advanced Dulbecco’s modified Eagle’s medium (DMEM) (1g/L glucose) with 10% (v/v) fetal bovine serum (FBS) and 1% (v/v) penicillin-streptomycin) and high glucose medium (high glucose Dulbecco’s modified Eagle’s medium (DMEM) (4.5 g/L glucose) with 10% (v/v) fetal bovine serum (FBS) and 1% (v/v) penicillin-streptomycin). To ensure that HaCaT cells were migrating and not proliferating during the experiments, they were exposed to mitomycin c (ThermoFisher Scientific) for 30 minutes. The cells were incubated for a total of 6 hours prior to commencing the actual experiments. 1μM of Oligomycin was added as a specific inhibitor to prevent the increase in mitochondrial respiration induced by ADP without inhibiting uncoupler stimulated respiration. All images were taken using an Olympus IX75 microscope with 5x, 10x, and 20x magnification.

### 2.3. Oxygen gradient application

After 6 h of culturing, cells in the microfluidics device were washed with DPBS and moved to an on-stage incubator on a microscope. Nitrogen and oxygen gas supplies were then connected to the device and adjusted to 18.3 and 7.14 sccm flow rates, respectively. The cells were incubated for 10 minutes to allow the gradient to stabilize, and then the experimental protocol described below was applied.

### 2.4. Oxygen measurements

The phase shift-based fiber optics fluorescence oxygen sensor (Ocean Optics, Dunedin, Florida) was used to characterize the gradients of the 11^th^ channels of the oxygen gradient device. To position the tip of the fiber over the desired position under the microscope, we used an XYZ micrometer that was mounted on a ring stand. The fiber tip had a 45° angle level and was pressed gently on the membrane to prevent tearing. It was important to ensure complete contact between the probe and the membrane to avoid fluctuations in measurements that could be caused by factors such as oxygen flux, fluid flow, and optical reflections from a surface. This was confirmed easily under the microscope when the tip was in focus. For measurements taken with HaCaT cells, the probe was positioned at the center of the gradient and the cells were removed from the PDMS surface before taking readings with the probe.

### 2.5. Migration quantification methods

The rate of cell migration can be quantified using a single metric or a combination of metrics. In this study, wound width and area, and wound closure percentage were calculated. Wound width was determined by measuring the distance between the edges of a scratch/stamp which decreases as cell migration progresses over time. The wound area was calculated by manually tracing the cell-free area in images using the ImageJ software. Normally, the wound area will decrease over time. Alternatively, the migration rate can be expressed as the percentage of area reduction or wound closure. The closure percentage increases as cells migrate over time. Grada et al. demonstrated how the scratch wound assay can be used to evaluate the migration capacity of keratinocytes in different experimental conditions (**Grada et al., 2017**).

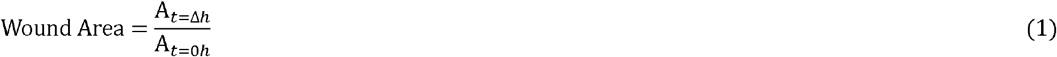

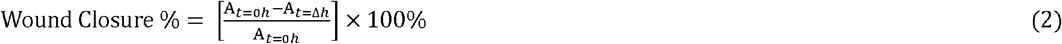

A_*t*=0*h*_ is the area of the wound measured immediately after scratching (t = 0h).

A_*t*=Δ*h*_ is the area of the wound measured h hours after the scratch is performed.

### 2.6. Mitochondrial and cytoplasmic ATP Imaging

ATP LW and red live cell fluorescence dye (BioTracker) were used to measure the ATP levels in the mitochondria and cytoplasm of HaCaT cells. After 5 h of incubation in the microfluidic device, the glucose free medium was removed and replaced with 2 mL low/high glucose medium containing 5 μM BioTracker dye. Cells were treated with streptomycin c as mentioned earlier to ensure the migration not proliferation of cells. The sample was then incubated for 15 minutes at 37°C and 5% CO_2_ before moving it to an on-stage incubator on a microscope. Cells in the gradient device were imaged for 24 hours.

### 2.7. Reactive oxygen species Quantification

Reactive oxygen species (ROS) were quantitatively and qualitatively determined using the cell permeant 2’,7’-dichlorodihydrofluorescein diacetate (H_2_DCFDA) (ThermoFisher Scientific). H2DCFDA is a type of fluorescein that has been chemically reduced and is used to detect the generation of reactive oxygen intermediates in neutrophils and macrophages. When the acetate groups are removed by intercellular esterases and oxidized, H2DCFDA changes from a non-fluorescent state to a highly fluorescent 2’,7’-dichlorofluorescein (DCF), which is used to indicate the presence of ROS in cells. Per the manufacturer’s instructions, the DCFDA was dissolved in DMSO to get 10 mM standard solution. The solution was then diluted 2 times to get a concentration of 10 μM. 1.2 μl of 10 μM solution was added to low/high glucose media and incubated for 24 h. 100 μL of the media was collected at 1, 3, 7, 12, 15, 20, and 24 h and were added to 96 well plate. The plate was then read using a BioTek Eon Microplate reader at 525 nm. The reactive oxygen species concentration in test samples was calculated using regression analysis. For qualitative analysis, samples were incubated for 24 hrs and images were then captured using an Olympus IX75 microscope with 10x magnification after 24 hours. 10 mM of Sodium Pyruvate was added to low/high glucose medium to inhibit the induced oxidative stress from hydrogen peroxide (H_2_O_2_). Fluorescence signal intensity was analyzed and quantified using ImageJ for normoxic and hypoxic gradient conditions.

### 2.8. Live/Dead assay

Cell viability was determined by a Live and Dead Cell Assay (Abcam). After HaCaT cells were attached to the device for 6 h, they were washed with PBS, and 5x diluted live and dead dye in PBS was added to the device. After 10 min of incubation, the device was put in on-stage incubator under a fluorescence microscope for 24 h. Images were taken via the automated X/Y stage to generate image collages. ImageJ software was then used to quantify and analyze the images. All measurements were conducted in triplicate. Cell viability was measured as a percentage using the following equation:

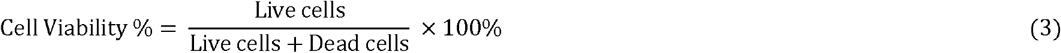

### 2.9. Statistical Analysis

All experiments were conducted at least in triplicates and repeated three times. The data represent the mean ± S.E.M. of three independent experiments. Statistical analysis was carried out using one-way ANOVA and regression analysis. The differences between the two sets of data were considered significant at p-value < 0.05.

## 3. Results

### 3.1. Oxygen microfluidic based cell migration

Coating PDMS with poly-L-lysine enhanced cell adhesion protein adsorption on the surface to promote cell migration. To visualize the gas microchannels, membrane cross sections were cut and imaged with brightfield microscopy as shown in Figure 2a. The images were captured using 5X, 10X, and 20X magnification to calculate the accuracy compared to the original model. An accuracy of 96.3% was achieved for microchannels of 376 μm width and 1000 μm thickness. To investigate the impact of varying levels of oxygen concentration on cell migration over the poly-L-lysine coated device, we applied the microfluidic oxygen gradients to HaCaT cells. Since the device is symmetrical, we captured one-half of the gradient through a 2 × 12 tiled microscopy collage at 10x magnification and then analyzed eleven positions within the 1-16% oxygen range (Figure 2b). To better check the dynamics for oxygen and nitrogen diffusion, an equilibration test was conducted using a fiberoptic probe over a period of 181s. The oxygen diffusion and nitrogen convection reached their equilibriums after 80 s (Figure 2c). The profile of oxygen across the channels shows concentrations from 1.62% to 15.69% (Figure 2d). We applied this calibration curve to present the oxygen gradient for the rest of the migration assays.

**Figure 2.**
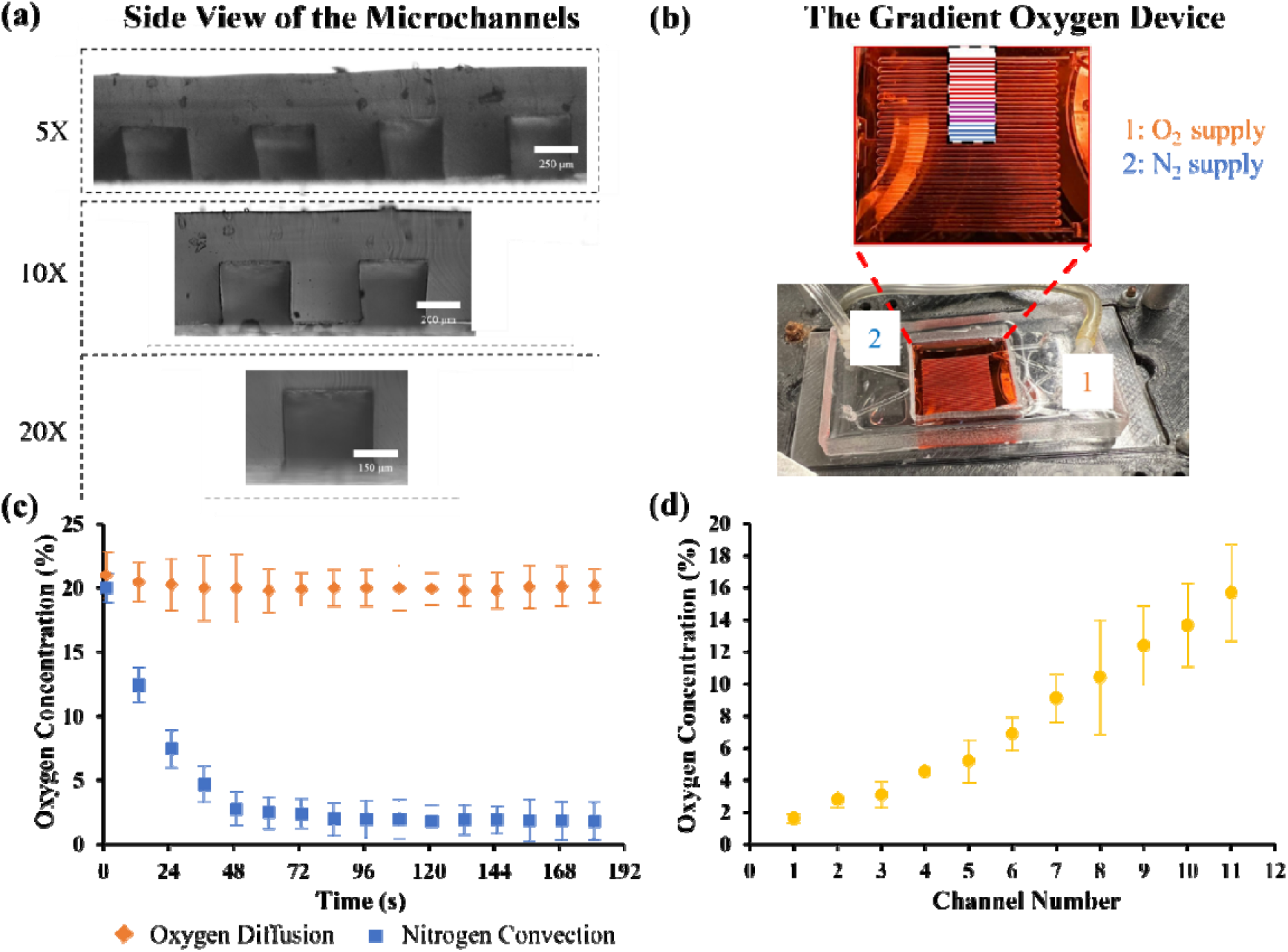
Oxygen gradient simulation with spatiotemporal detection. (a) Images showing the 376 μm microchannels width with different magnifications (5X, 10X, and 20X). They showed an accuracy of 96.3% compared to the original model. (b) Eleven positions with the oxygen gradient were selected as regions of interest for cell migration and mitochondrial and cytoplasmic ATP responses. (c) Oxygen concentration was quantified using a fiber optics fluorescence oxygen sensor. The measurements were taken at the diffusion spots of the gradient oxygen device for 181s with a time interval of 12s. (d) Oxygen concentration was measured at the middle of each channel along the eleventh channel of the device. O_2_ concentration was linearly increasing between channels 1 and 4 and then exponentially jumped to reach 15.69% at channel 11. Error bar S.E.M (N=3).

### 3.2. Hypoxic gradient enhanced cell migration at low glucose

Stamp and scratch wound assays were carried out to determine the effect of low and high glucose incubations on subsequent migration of HaCaT cells with (hypoxic gradient) and without (normoxia) oxygen gradients. For both low and high-glucose cultures, the stamp assay showed less average migration compared to scratch assays (Figure 3a). For stamp assays, migration was higher in low versus high-glucose cultures. Additionally, the incorporation of the hypoxic gradient advanced the average migration (over the whole gradient) for both glucose conditions (Figure 3b). For example, for low glucose normoxic conditions, the migration distance after 24 h had a range between 65.00 ± 5.80 μm and 130.00 ± 11.20 μm which is slightly higher than that of high glucose condition of a range between 35.00 ± 7.30 μm and 70.00 ± 12.60 μm (Figure 3b). Further, it was observed that the migration distance for hypoxic gradient conditions of low glucose had a range between 95.00 ± 13.20 μm and 170.00 ± 20.40 μm which is higher than both, high glucose conditions (65.00 ± 6.10 μm and 110.00 ± 11.60 μm) and low glucose of normoxic conditions (Figure 3b).

**Figure 3.**
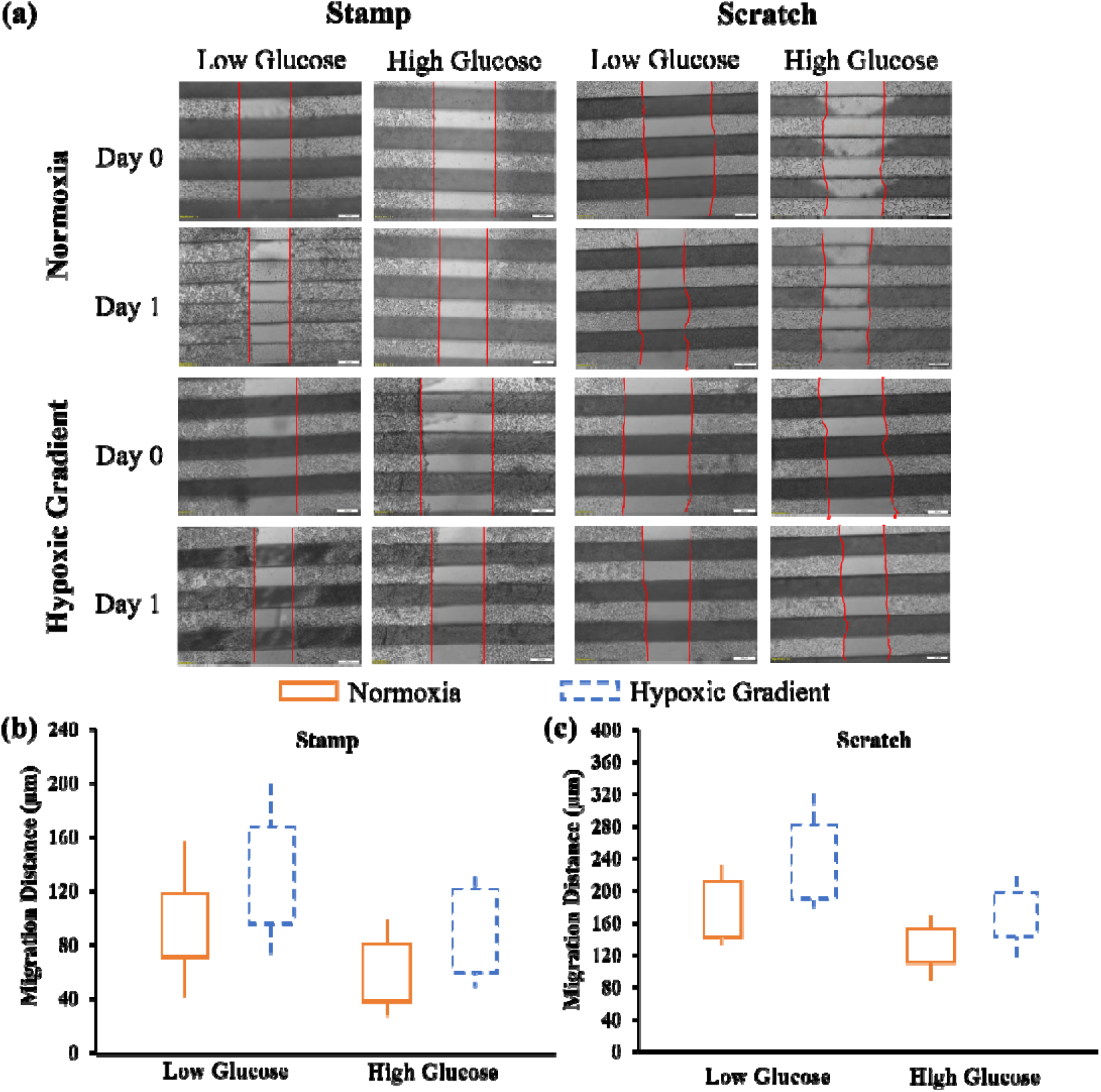
Oxygen gradient stimulates cell migration in HaCaT cells during wound healing. (a) Images of cell migration were captured using an Olympus IX75 microscope with 20x magnification at the time of scratch/stamp and 24 hours after to measure the migration distance, wound area, and wound closure (%). For both (b) stamp and (c) scratch assays, the migration distance was measured using ImageJ software for low and high glucose conditions with and without the oxygen gradient. The average oxygen gradient is the average of the oxygen concentrations all over the eleven channels.

Similarly, for scratch assays, migrations were faster for low glucose of hypoxic gradient conditions over their counterparts (Figure 3c). The migration distance for low glucose conditions showed an increase from 145.00 ± 23.80 μm and 210.00 ± 30.50 μm for normoxic conditions to 187.00 ± 9.40 μm and 280.00 ± 37.10 μm for hypoxic gradient conditions. The same trend was shown for high glucose with a slight increase from 115.00 ± 18.30 μm and 150.00 ± 22.30 μm of normoxic conditions to 140.00 ± 20.90 μm and 195.00 ± 27.20 μm of hypoxic gradient conditions.

These results suggest that while high glucose impairs keratinocyte migration, hypoxia stimulation evidently enhances it, with the oxygen gradient showing a more prominent impact on scratch (injury dependent) assays (Figure 3c). Therefore, oxygen gradient improved the migration of HaCaT cells in both glucose conditions, especially for scratch assay. On the contrary, high glucose conditions inhibited the migration of cells while low glucose enhanced keratinocyte migration in hypoxic conditions.

To further describe the migration of HaCaT cells under the hypoxic gradient, the wound area between all eleven oxygen concentrations was analyzed (Figure 4a). For stamp assays, results showed more wound closure (smaller area) for low glucose conditions than for high glucose conditions across the gradient. The final wound area ranged from 0.49 ± 0.03a.u. to 0.56 ± 0.01a.u. versus 0.81 ± 0.02a.u. to 0.87 ± 0.01a.u. for low versus high glucose stamp assays. Little to no difference was observed across gradient concentrations for stamp high glucose condition. On the other hand, the scratch assay showed better overall wound closure than stamp assays with a similar trend of low glucose yielding smaller final wound area than high glucose. The final wound areas ranged from 0.41 ± 0.01a.u. to 0.48 ± 0.02a.u. versus 0.55 ± 0.03a.u. to 0.62 ± 0.01a.u. for low versus high glucose scratch assays. Furthermore, significant modulation was observed around channel 7 in the gradient, corresponding to an oxygen concentration of ∼9%. For comparison, the wound area was averaged across the whole chamber for low and high glucose conditions with and without the hypoxic gradients. Results indicated that the hypoxia significantly increased the migration of HaCaT cells (Figure 4b). The scratch assay had a decrease in wound area from 53.00 ± 2.81% to 41.00 ± 1.90% and 77.00 ± 5.43% to 55.00 ± 3.10% (p-value <0.001) for low and high glucose conditions between normoxia and hypoxic gradient, respectively. Similarly, the stamp assay had a decrease in wound closure from 76.19 ± 5.20% to 49.00 ± 2.70% and 94.72 ± 9.10% to 81.00 ± 7.40% for both conditions (p-value<0.001). These results suggest that while glucose impaired cell migration in both assays, injury in the scratch assay initiated an additional oxygen modulation that is not observed in the stamp assay without injury.

**Figure 4.**
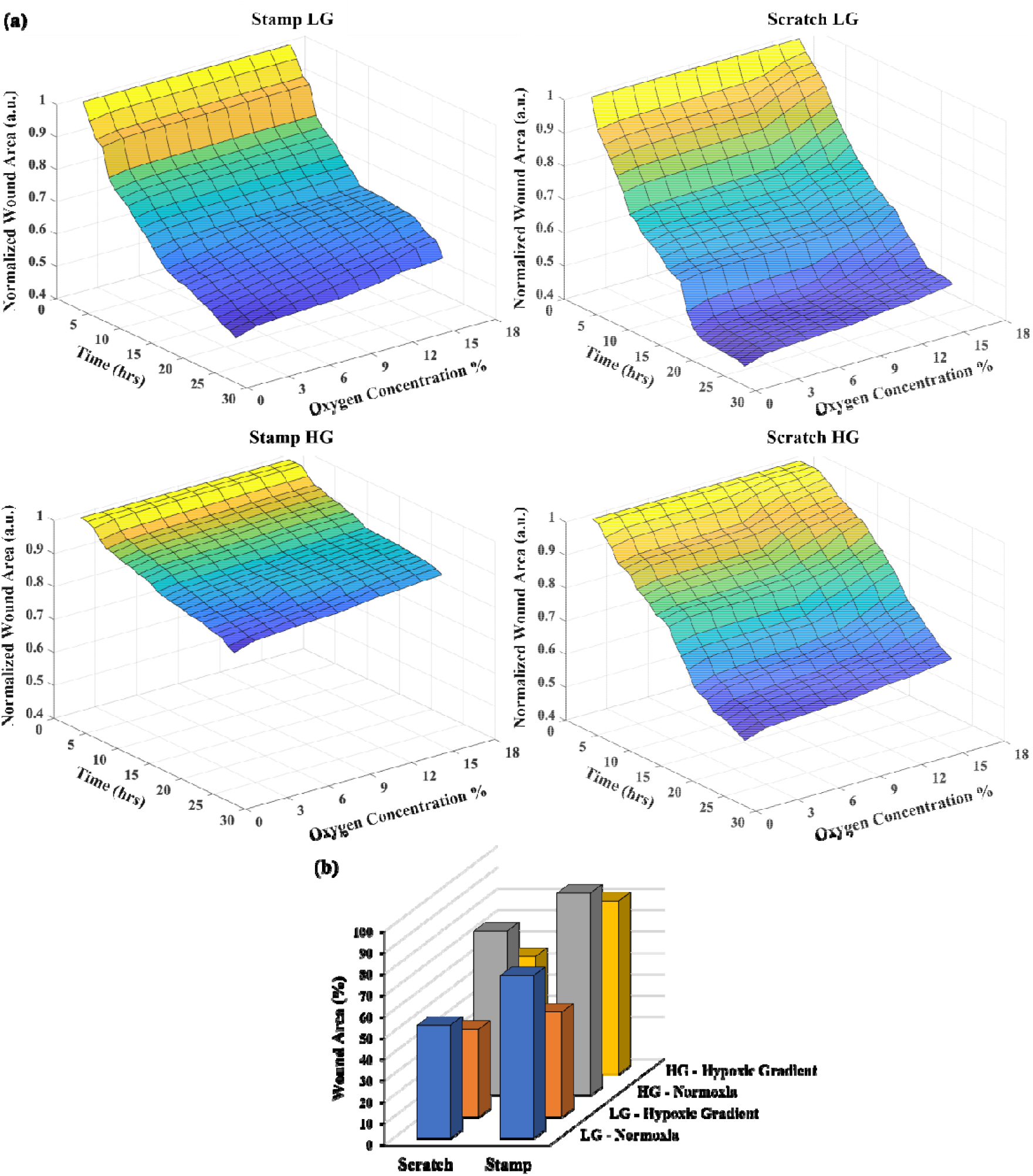
The hypoxic gradient promoted wound healing. (a) The wound area was measured for all channels in the cell migration experiment of HaCaT cells, with the wound area computed for each of the 11 channels and normalized to the wound at time 0. Results indicated that the wound area remained relatively high until the 12^th^ hour of the experiment for low glucose conditions in both the stamp and scratch assays. However, for high glucose, the wound area remained high throughout the 24 hours in the stamp assay while it decreased in the scratch assay, reaching almost 0.59 ± 0.02 at the 24^th^ hour. Additionally, (b) the wound area (%) was calculated for both conditions and assays, without (Normoxia) and with (Hypoxic gradient) the oxygen gradient. Results showed that the oxygen gradient significantly promoted the migration of HaCaT cells.

### 3.3. A tradeoff between mitochondrial versus cytoplasmic ATP

To better assess how consistent our gradient oxygen device is, control experiments were run for mitochondrial and cytoplasmic ATP for low and high glucose conditions. Results showed that both the device and the petri dish had almost the same ATP intensity (Figures 5a, b, d, and e) with a difference of less than 10% for all conditions (Figure 5c).

**Figure 5.**
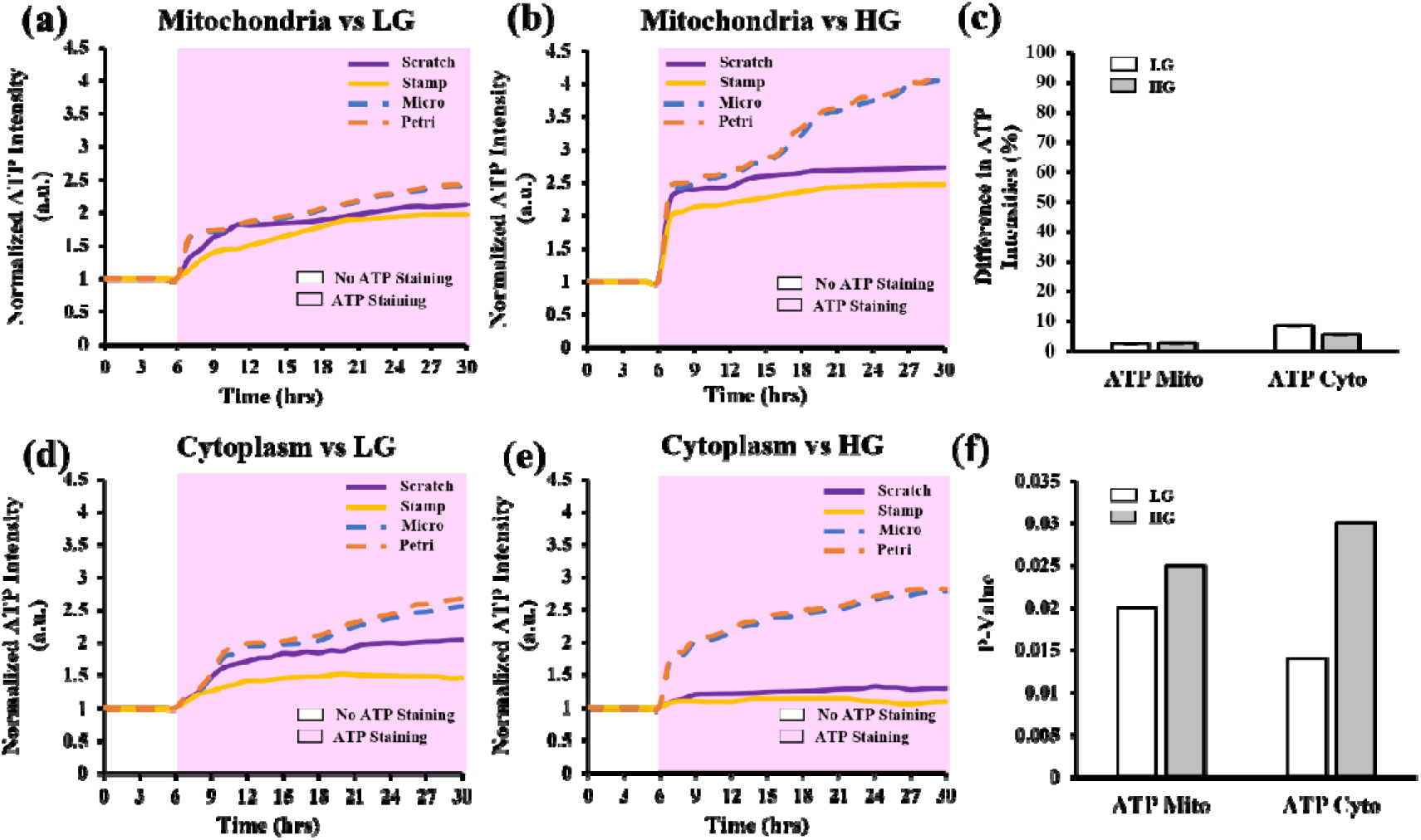
ATP fluorescence detection without gradient oxygen stimulation (Normoxia). The intensities of mitochondrial ATP were measured for (a) LG and (b) HG, and their respective intensity differences after 24 hrs were recorded as (c). The same study was conducted for cytoplasmic ATP, measuring (d) LG and (e) HG intensities and recording the intensity difference as (c). The experiments also included the representation of (f) P-values for all the measurements. High glucose (HG) conditions elevated the mitochondrial ATP for scratch and stamp with almost no cytoplasmic ATP, while low glucose (LG) had lower mitochondrial ATP but higher cytoplasmic ATP for both assays. These results showed that HaCaT cells with scratch low glucose condition had higher cytoplasmic ATP indicating active metabolism and cell migration.

We further measured the levels of mitochondrial and cytoplasmic ATP to gain a deeper understanding of how the migration of HaCaT cells is affected by glucose and hypoxic conditions in the gradient device. Under low glucose, scratch assay HaCaTs had higher mitochondrial and cytoplasmic ATP levels (Figures 5a, d) compared to stamp (P-value<0.05). With high glucose culturing, the ATP levels were relatively the same between scratch versus stamp assays (P-value<0.05). This suggested that cells under injury in scratch assays were metabolically more active and made use of the excess glucose exposed to them. Additionally, there is a trade-off between mitochondrial versus cytoplasmic ATP levels under different glucose culturing conditions. High glucose culturing has elevated mitochondrial ATP compared to low glucose cells (Figures 5b, e). However, this trend is flipped on the cytoplasmic side, where low glucose cells have more ATPs than high glucose ones (P-value<0.05). We suspect that high glucose is diverting the ATP metabolism away from the cytoplasm, reducing the available energy for changes in the cytoskeleton required for cell migration.

To further investigate the mitochondrial and cytoplasmic ATP production under a hypoxic gradient, ATP fluorescence images for stamp and scratch assays were quantified using ImageJ. For stamp assays, Figure 6a, the same switch from cytoplasmic to mitochondrial ATP was seen when going from low to high glucose—mitochondrial ATP rose, and cytoplasmic ATP fell across the gradient. However, there are distinct dynamics near 4.16-9.14% hypoxia in low glucose cells, where mitochondria ATP dipped, and cytoplasmic ATP peaked compared to other oxygen levels. These dynamics were less obvious in the high glucose cells, which already had elevated mitochondrial and minimal cytoplasmic ATP levels. This suggests that for injury-free stamp migration, specific levels of hypoxia—i.e. 4.16-9.14%—promote higher cytoplasmic ATP that can be utilized in cell migration, especially with low glucose conditions. When cultured in high glucose, however, this hypoxic promotion is overwhelmed by a switch to predominantly mitochondrial metabolism.

**Figure 6.**
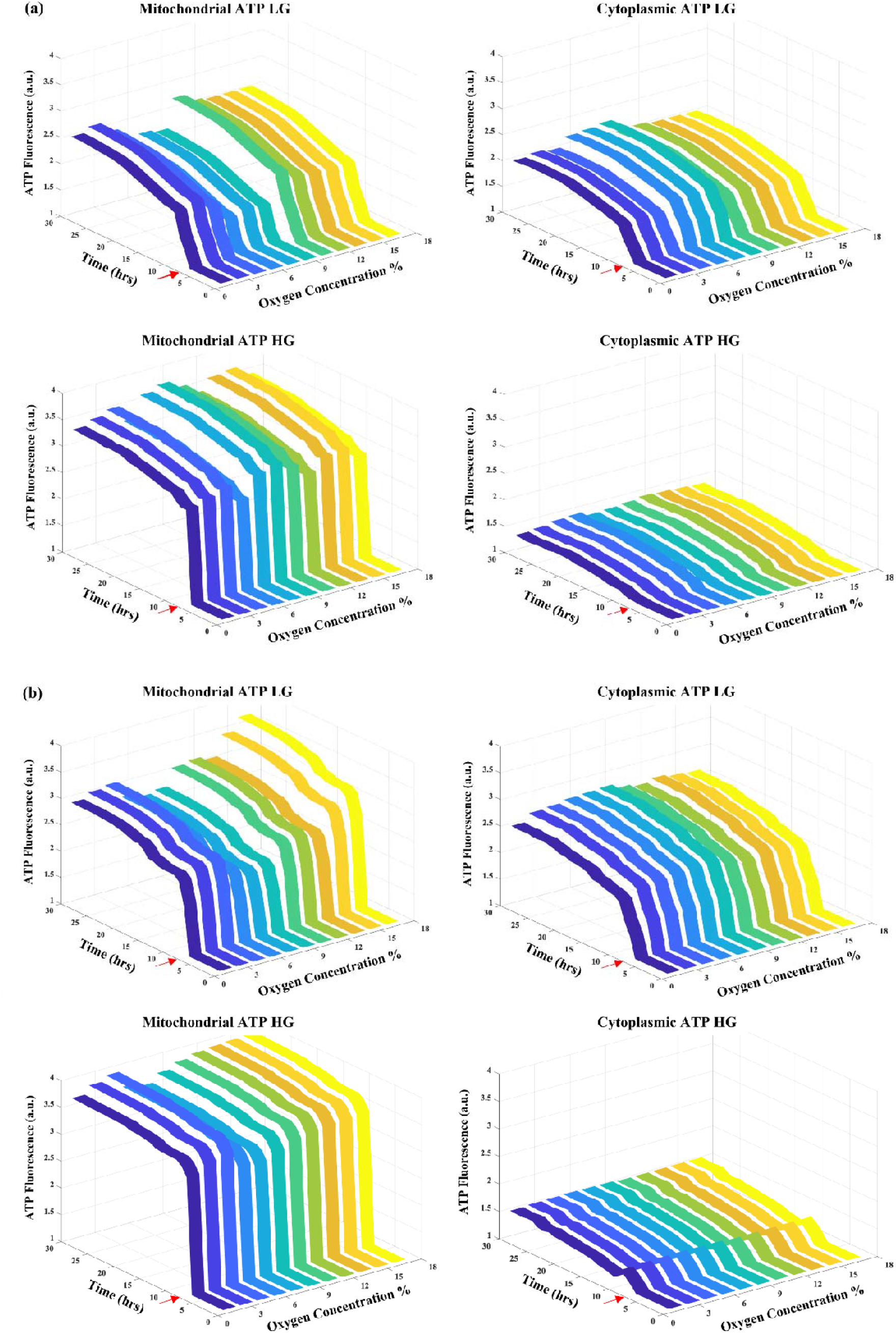
Oxygen gradient stimulation with ATP fluorescence detection for stamp and scratch assays. Mitochondrial and cytoplasmic ATP with gradient for (a) stamp and (b) scratch was quantified via fluorescence labeling, showing modest increases until the same 4.16-9.14% oxygen range, where the response dipped before rising again near 18.00%. These results suggested that HaCaT cells didn’t mount ATP responses to high glucose. The red arrow shows the beginning of the ATP staining time after culturing cells for 6 hrs.

When an injury is introduced via the scratch assay, we see similar ATP trends presented in the stamp assay with one distinct difference under high glucose (Figure 6b). In general, low glucose had higher cytoplasmic ATP levels while high glucose had higher mitochondrial levels. The same trade-off dynamics near 4.16-9.14% hypoxia was present in low glucose scratch assay, albeit not as obvious with almost 2.46 ± 0.21a.u. for mitochondrial ATP and 2.47 ± 0.14 a.u. for cytoplasmic ATP. However, while this dynamic was not seen in high glucose stamp assays, the dip in mitochondrial ATP was preserved in the high glucose scratch assay. Furthermore, all ATP levels from scratch assays were higher in general when compared to stamp assays. This result suggests that when there is injury introduced in the model, HaCaT cells can leverage more ATP metabolism for migration with a preference for certain levels of hypoxia. And this hypoxic sensitivity is maintained even in high glucose levels compared to the stamp assay. To put these findings in the scope of diabetic wound healing, glucose alters ATP metabolism away from cell migration and hypoxic signaling, but injury recovers this hypoxic signaling for cell migration.

### 3.4. Reactive oxygen species modulate the migration of HaCaT cells under high glucose

Reactive oxygen species (ROS) are highly reactive molecules that can cause damage to cellular components, such as DNA, proteins, and lipids. In excess, ROS can lead to oxidative stress, which can be harmful to cells and contribute to various diseases (**Milkovic et al., 2019**). In the context of cell migration, studies have shown that high levels of ROS can inhibit cell migration by interfering with the cytoskeletal rearrangements necessary for the process. ROS can cause the oxidation of actin, which is a major component of the cytoskeleton, leading to its depolymerization and inhibiting the formation of lamellipodia and filopodia which are essential for cell migration. Additionally, ROS can activate signaling pathways that negatively regulate cell migration, such as the RhoA/ROCK pathway, which can induce the formation of stress fibers and inhibit cell motility (**Zi et al., 2018**). Therefore, ROS can act as a negative regulator of cell migration, and the balance between ROS production and scavenging is crucial for maintaining proper cellular functions.

The aim of this experiment was to measure the fluorescence signal intensity of reactive oxygen species via quantitative and qualitative analysis for normoxic and hypoxic gradient conditions for stamp and scratch assays. ROS intensity of high glucose conditions was the highest compared to low glucose conditions of a scratch assay for normoxic and hypoxic conditions (Figures 7a, b, and c). The same trend was observed for stamp assay (Figures 7a, b, and c) but with higher intensities. For normoxic conditions, the normalized extracellular ROS intensity showed an exponential increase from 1.10 ± 0.01a.u. at hour 6 to 2.08 ± 0.02 a.u. at hour 18, followed by a jump to reach 2.70 ± 0.02 a.u. at hour 21 then a steady state until hour 26, and then a further increase to 3.10 ± 0.01 a.u. at hour 30 for stamp high glucose (Figure 7d). A similar pattern was observed for high glucose conditions of the scratch assay with a linear increase from 1.06 ± 0.02 a.u. at hour 6 to 2.69 ± 0.01 a.u at hour 30 with normoxia (Figure 7d). However, a different trend was observed for low glucose conditions with normalized intensity near 1 for scratch and stamp assays with normoxia (Figure 7d). In contrast, under hypoxic gradient conditions, both assays showed a similar trend for both low and high glucose conditions, with a slightly higher signal compared to normoxic conditions (Figure 7e). These results indicate that irrespective of the assays, the signal intensity was the highest for high glucose conditions which inhibited the migration of HaCaT cells.

**Figure 7.**
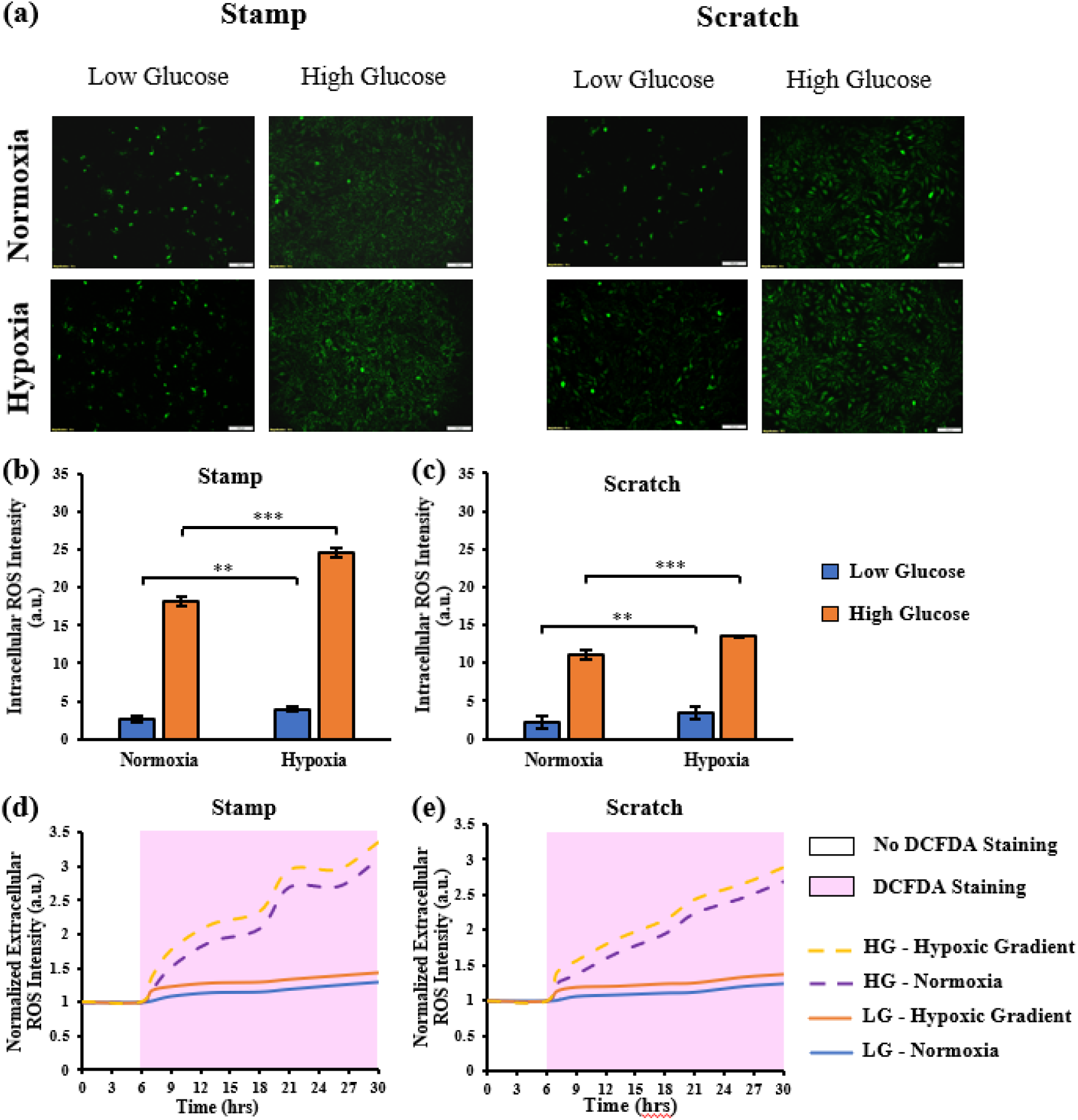
Detection of reactive oxygen species (ROS) via fluorescence and microplate reader intensity for normoxic and hypoxic gradient conditions. (a) Images of ROS were captured using an Olympus IX75 microscope with 10x magnification after 24 hours. DCFDA reagent was used to detect the reactive oxygen species. Fluorescence signal intensity was analyzed using ImageJ for (b) stamp and (c) scratch assays. High glucose in both conditions had the highest fluorescence with a slightly higher signal for hypoxic gradient than in normoxic conditions. The fluorescence signal from the microplate reader was also measured and normalized to the values collected after 30 minutes of starting the experiment for (d) stamp and (e) scratch assays. The same trend was found compared to the data collected from fluorescence signal intensities for both conditions. Error bar S.E.M (N=3). **p-value <0.01 and ***p-value <0.001.

### 3.5. Role of sodium pyruvate and oligomycin in mitigating ATP imbalances

Elevated levels of glucose have demonstrated a detrimental effect on the movement of HaCaT cells, resulting in increased release of reactive oxygen species (ROS) and consequent inhibition of cell migration. To restore proper metabolic states for the migration process, 1μM oligomycin was added to modulate lactic acid and block aerobic respiration, and 10mM sodium pyruvate was added as a ROS scavenger. To gain a more comprehensive understanding of cell behavior under these circumstances, the intensities of mitochondrial and cytoplasmic ATP were measured over a span of 24 hours, both under normoxic and hypoxic conditions.

Under normoxic conditions, sodium pyruvate exhibited a slight impact on mitochondrial and cytoplasmic ATP levels for stamp assay (Figures 8a, d). However, in the context of scratch assays conducted under normoxic conditions, sodium pyruvate had a greater influence, lowering mitochondrial ATP levels while elevating cytoplasmic ATP levels (Figures 8b, e). Oligomycin had a significant effect on HaCaT cell metabolism, particularly in the case of scratch assays performed under normoxic conditions (Figures 8b, e). By the 24^th^ hr, the normalized mitochondrial ATP intensity for scratch assays reached a value of 1.79 ± 0.02 a.u., which was lower than the corresponding values of 2.46 ± 0.01a.u. and 2.73 ± 0.03 a.u. observed with the addition of 10mM sodium pyruvate and the control condition (high glucose only), respectively (Figures 8b, c) (p-value<0.05). As for cytoplasmic ATP, the trend was reversed, with scratch assays resulting in a level of 1.81 ± 0.01 a.u. when 1μM oligomycin was added, compared to 1.37 ± 0.02 a.u. with 10mM sodium pyruvate and 1.29 ± 0.01 a.u for the control (Figures 8e, f) (p-value<0.05). In the case of scratch assays performed under normoxic conditions, the combined presence of sodium pyruvate and oligomycin led to lower mitochondrial ATP levels and higher cytoplasmic ATP levels. The same trend was observed under hypoxic conditions but with higher ATP levels compared to normoxia in both assay types (Figure 9) (p-value<0.01). These findings indicate that the inclusion of 1μM oligomycin and 10mM sodium pyruvate effectively restored the migration of HaCaT cells under high glucose conditions, particularly when an oxygen gradient was applied.

**Figure 8.**
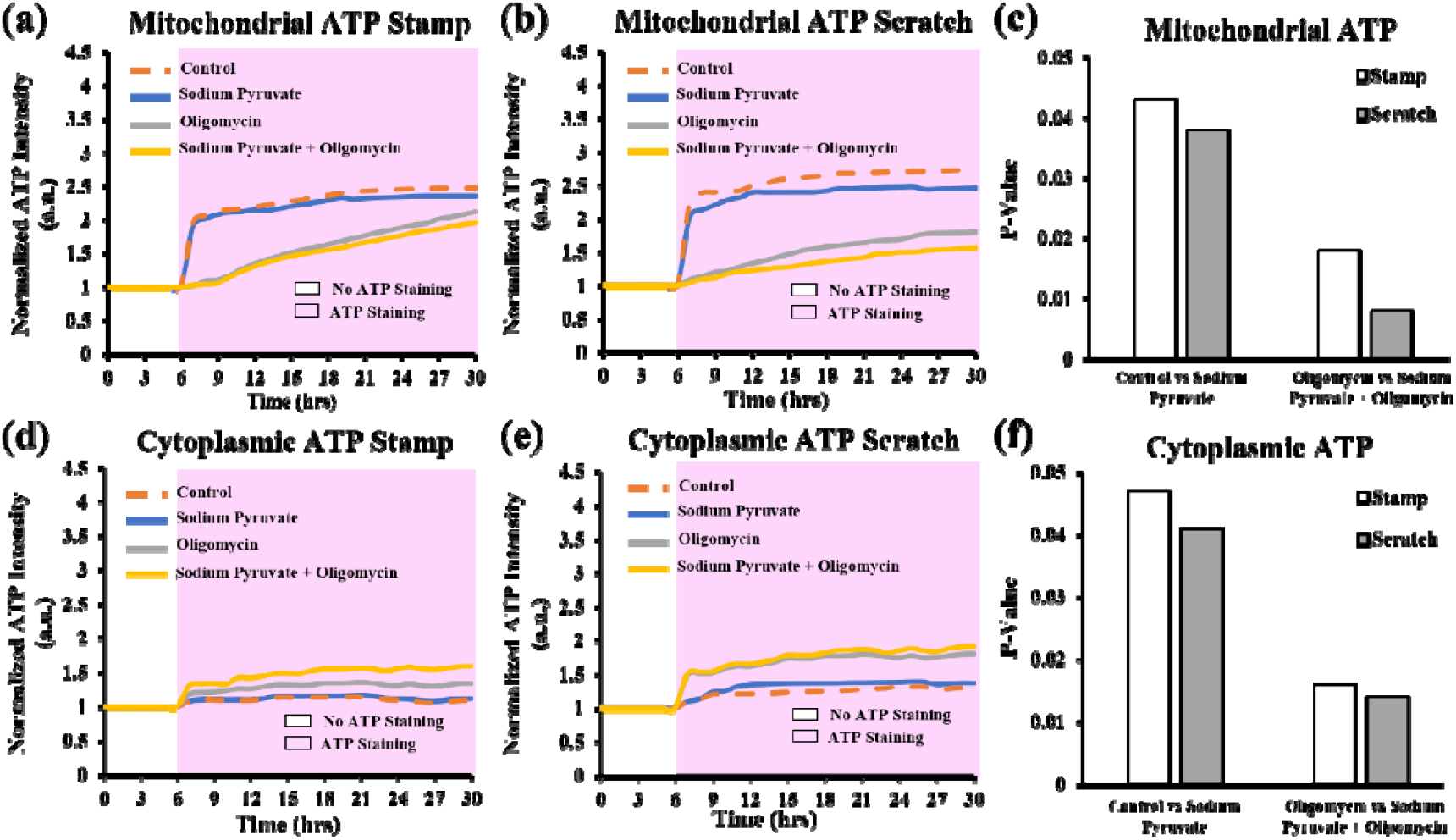
ATP fluorescence detection without gradient oxygen stimulation (Normoxia) with ROS scavenger and lactic acid. The intensities of mitochondrial ATP were measured for (a) stamp and (b) scratch, and the p-values were calculated and shown as (c). The same study was conducted for cytoplasmic ATP, measuring (d) stamp and (e) scratch ATP intensities and showing p-values as (f). 10mM sodium pyruvate and 1μM oligomycin elevated the cytoplasmic ATP levels for scratch with normoxic conditions.

**Figure 9.**
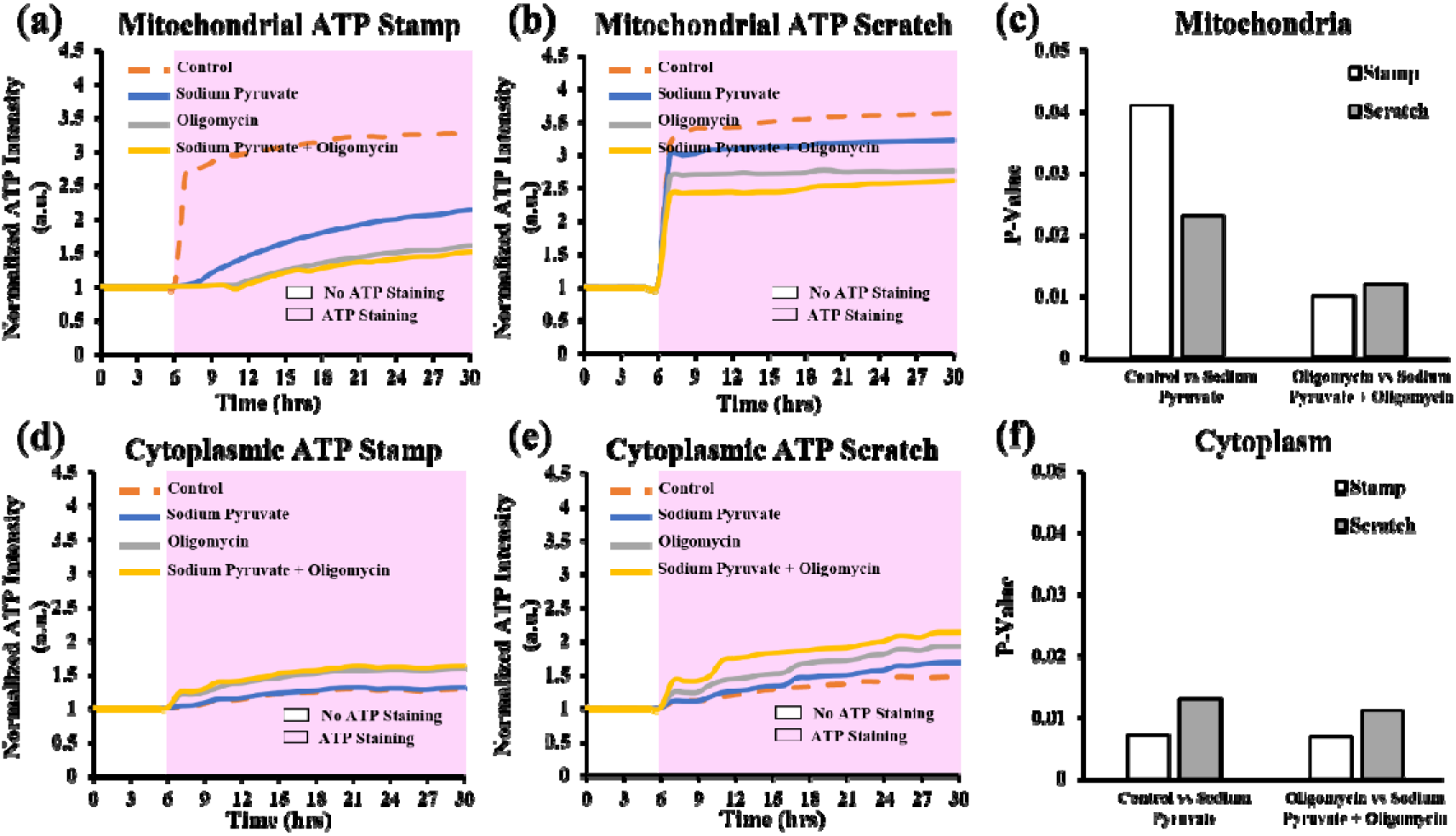
ATP fluorescence detection with gradient oxygen stimulation (Hypoxic gradient) with ROS scavenger and lactic acid. The intensities of mitochondrial ATP were measured for (a) stamp and (b) scratch, and the p-values were calculated and shown as (c). The same study was conducted for cytoplasmic ATP, measuring (d) stamp and (e) scratch ATP intensities and showing p-values as (f). 10mM sodium pyruvate and 1μM oligomycin elevated the cytoplasmic ATP levels for scratch under hypoxic conditions. This effect is greater under hypoxia than that under normoxia (p-value<0.01).

### 3.6. Modulation of reactive oxygen species levels and HaCaT cell migration by oligomycin and sodium pyruvate in normoxic and hypoxic environments

The impact of oligomycin and sodium pyruvate on the intensity of the fluorescence signal produced by reactive oxygen species was measured using image J. Under normoxic conditions with stamp assay, the intracellular intensity of reactive oxygen species (ROS) decreased from 18.12 ± 1.06 a.u. (control as high glucose only) to 8.23 ± 0.80 a.u. with sodium pyruvate, 7.92 ± 0.71 a.u with oligomycin, and 7.33 ± 0.65 a.u with sodium pyruvate and oligomycin (Figures 10a, b). However, under hypoxic conditions with stamp, the ROS intensity decreased even further from the control level of 24.6 ± 1.43 a.u to 10.94 ± 1.13 a.u with sodium pyruvate, 8.56 ± 0.83 a.u. with oligomycin, and 8.11 ± 0.79 a.u. with sodium pyruvate and oligomycin (Figures 10a, b). These findings indicate that sodium pyruvate effectively scavenged the reactive oxygen species in the stamp assay, particularly in hypoxic conditions. The same trend was observed in the scratch assay, although the difference in ROS intensity between the control and other conditions was smaller (Figures 10a, c, d). This can be attributed to the lower ROS intensity observed in HaCaT cells during the scratch assay, regardless of normoxic or hypoxic conditions without oligomycin and sodium pyruvate. Therefore, the combination of the oxygen gradient and sodium pyruvate and oligomycin successfully reduced the ROS intensity in all conditions, especially in the scratch assay, which involves an actual injury.

**Figure 10.**
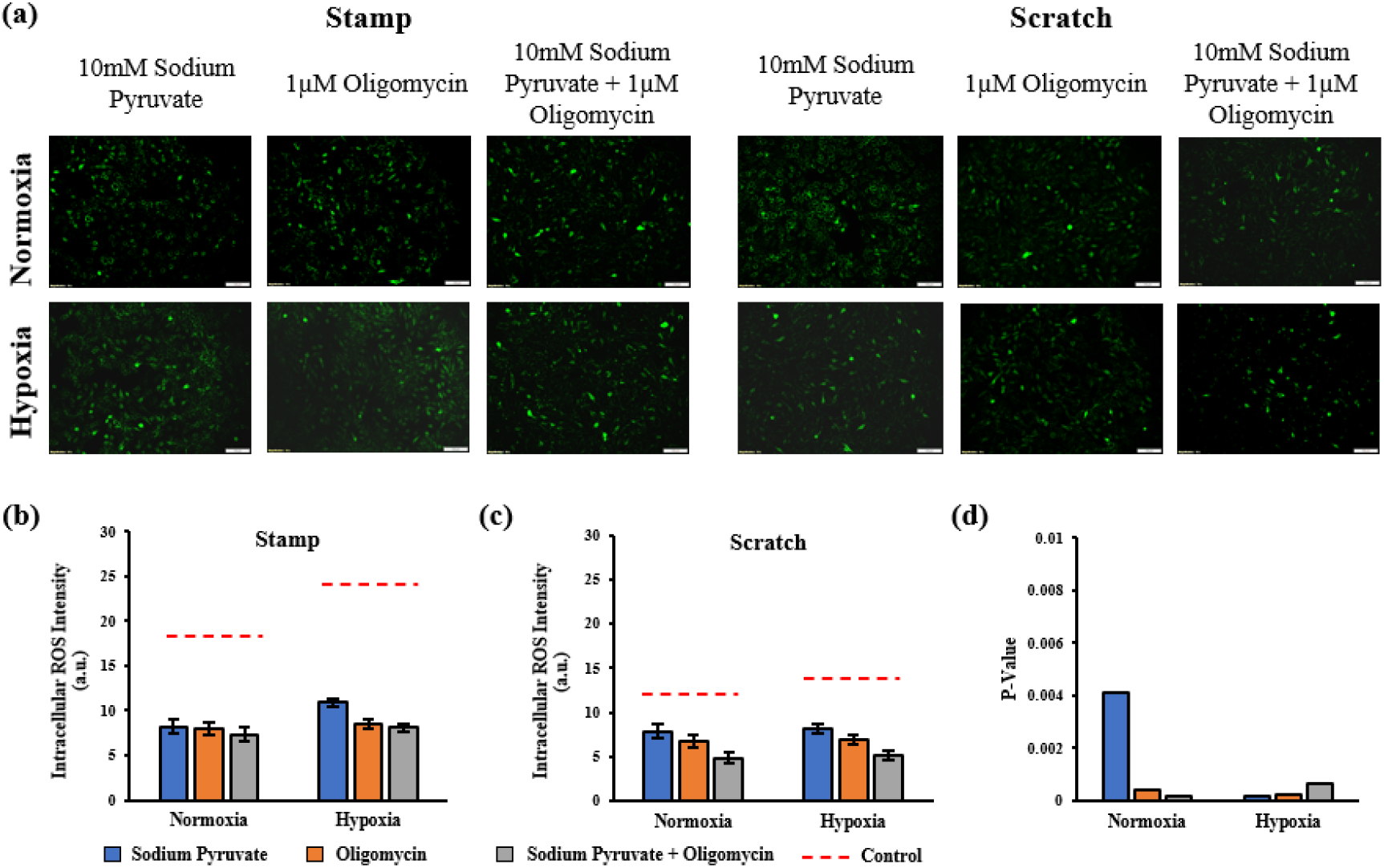
10mM sodium pyruvate and 1μM oligomycin lowered the reactive oxygen species expression for high glucose normoxic and hypoxic gradient conditions. (a) Images of ROS were captured using an Olympus IX75 microscope with 10x magnification after 24 hours. Fluorescence signal intensity was analyzed using ImageJ for high glucose conditions with 10mM sodium pyruvate and 1μM oligomycin for (b) stamp and (c) scratch assays. (d) p-values of the stamp-scratch assays for normoxic and hypoxic conditions.

Furthermore, the effect of adding 10mM sodium pyruvate and 1μM oligomycin on the migration of HaCaT cells under high glucose levels was assessed in both stamp and scratch assays using image J. The migration distance was found to increase from 35.00 ± 7.30 μm and 70.00 ± 12.60 μm in the stamp control to 80.00 ± 6.80 μm and 122.00 ± 13.10 μm in the stamp assay with sodium pyruvate and oligomycin under normoxic conditions (Figures 11a, b) (p-value<0.001). In hypoxic conditions, the migration distance in the stamp assay increased from 65.00 ± 6.10 μm and 110.00 ± 11.60 μm to 120.00 ± 15.21 μm and 170.00 ± 12.37 μm with sodium pyruvate and oligomycin (p-value<0.001). The same pattern was observed in the scratch assay, where the highest migration distance ranged between 180.00 ± 16.74 and 230.00 ± 21.12 when using 10mM sodium pyruvate and 1 μM oligomycin (Figures 11a, b) (p-value<0.001). Under normoxic conditions, when oligomycin and sodium pyruvate were introduced, the wound area decreased from 94.72 ± 3.71% to 79.16 ± 1.38% for the stamp assay, and from 77.00 ± 1.23% to 71.00 ± 1.74% for the scratch assay (Figure 11c) (p-value<0.001). Similarly, under hypoxic conditions, the wound area decreased from 81.00 ± 2.83% to 68.00 ± 1.32% for the stamp assay, and from 55.00 ± 0.97% to 51.00 ± 1.02% for the scratch assay (Figure 11c) (p-value<0.001). These results indicated that oligomycin inhibited mitochondrial ATP synthesis, while the addition of sodium pyruvate provided antioxidant support by scavenging reactive oxygen species (ROS) at the side of oxidative stress production. This antioxidant activity helped preserve mitochondrial function and integrity, ultimately regulating cell metabolism and facilitating cytoplasmic ATP production.

**Figure 11.**
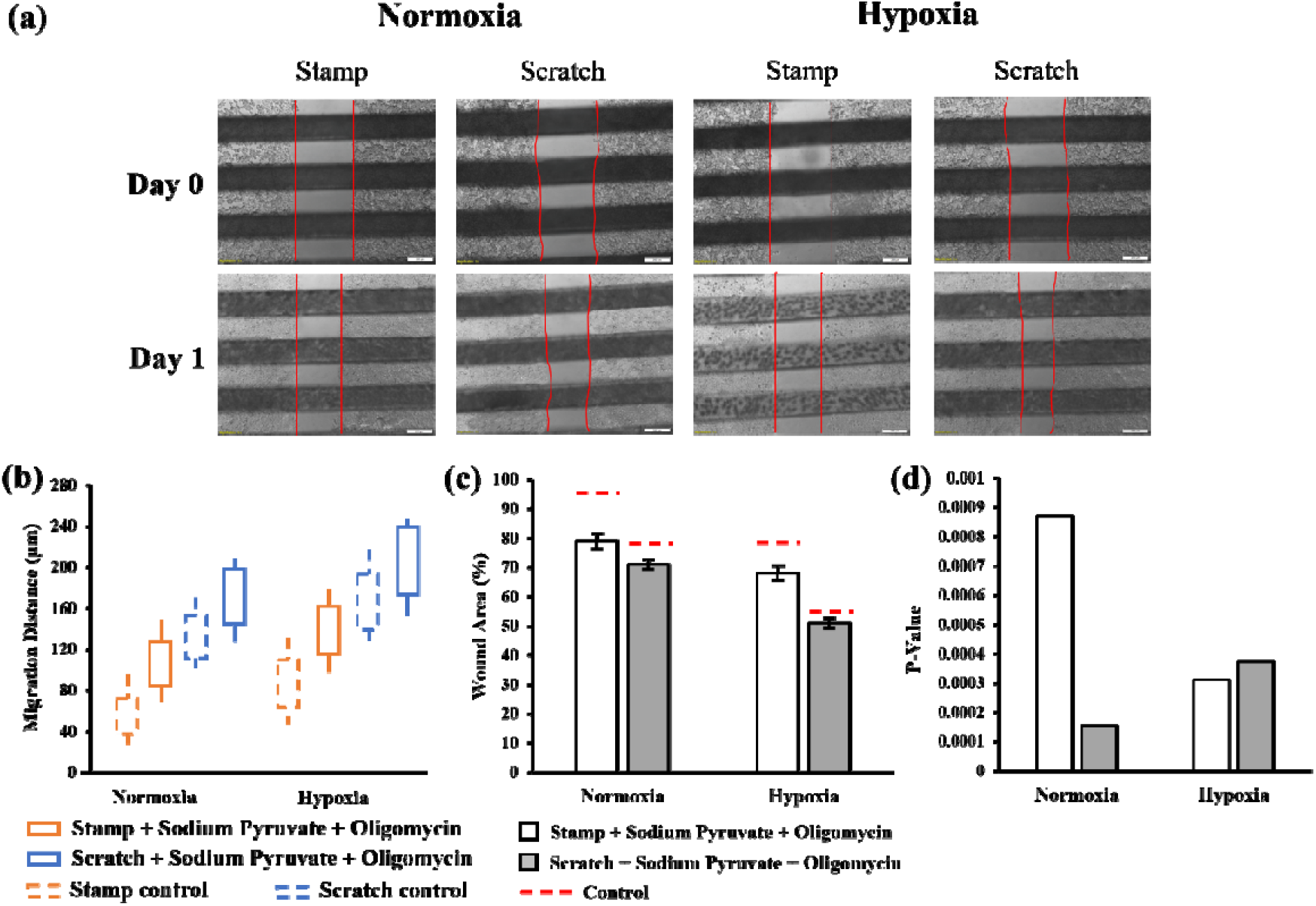
10mM sodium pyruvate and 1μM oligomycin rescued HaCaT cell migration for high glucose normoxic and hypoxic gradient conditions. (a) Images of cell migration were captured using the same microscope with 20x magnification at the time of scratch/stamp and after 24 hours. For both (b) migration distance and (c) wound area, ImageJ software was also used to take the measurements for high glucose conditions of stamp and scratch assays with and without the average oxygen gradient. (d) p-values of the control-stamp and control-scratch wound area analysis for normoxic and hypoxic conditions with 10mM sodium pyruvate and 1μM oligomycin.

## 4. Discussion

Microfluidic devices have become popular in disease modeling due to their ability to provide precise and controlled microenvironments for cell cultures (**Orabi et al., 2023**). Here we studied the impact of hyperglycemia and hypoxia on HaCaT cell migration using a gas microfluidic gradient with oxygen concentrations from 1.62% to 15.69%. Cell migration is fundamental in embryonic development, wound healing, metastasis, and other processes. The migration of HaCaT cells is initiated by various signals, including chemical and physical stimuli, that induce changes in cell behavior (**Veeraperumal et al., 2020; Zhou et al., 2023**). These stimuli can activate signaling pathways that lead to cytoskeletal remodeling, formation of focal adhesions, and secretion of proteases, all of which contribute to cell movement (**Qin et al., 2018; D’Aiuto et al., 2022**). Oxygen and metabolic gradients are two specific stimuli that are altered in diabetes.

First, our results suggested that a hypoxic gradient provides a cue to enhance migration. More specifically, 9% oxygen was a transition level that further advanced migration. This level of oxygen is neither hypoxic nor normoxic in a typical cell culture sense. However, we have also seen similar transition oxygen levels for other cell types such as epithelial and pancreatic islet cells (**Zeitouni et al., 2016**). This fits the unique tissue oxygen levels described by researchers in tissue hypoxia (**Komatsu et al., 2018**). Although hypoxia is well-studied for angiogenesis, regeneration, differentiation, and cell survival **(Lv et al., 2017; Okonkwo and DiPietro, 2017)**, its role in wound healing is still unclear. When exacerbated by systemic diseases like diabetes, the hypoxia-hyperglycemia mechanism becomes further complicated. Furthermore, high glucose incubation slowed migration and reduced the effects of the aforementioned hypoxia cues. For injury-free stamp assays, low glucose results showed moderate hypoxic modulation, whereas high glucose was devoid of any modulation across the gradient. On the other hand, injury provided in the scratch assays recovered this hypoxic sensitivity. This suggested that migration depends on a combination of hypoxic and metabolic cues presented by injured cells in the wound bed. These findings prompted us to examine ATP metabolism in more detail.

Much of the cytoskeletal and focal adhesion mechanisms in HaCaT migration are dependent on ATP metabolism. Because the production and utilization of ATP are localized differently, we examined the expressions of mitochondrial versus cytoplasmic ATP during stimulated HaCaT migration. Mitochondrial ATP generated through oxidative phosphorylation requires oxygen to complete the electron transport chain **(Townsend et al., 2020; Boyman et al., 2020)**. In contrast, cytoplasmic ATP is generated mainly through glycolysis. Additionally, under conditions of low oxygen availability, such as during intense exercise, ATP can also be generated through anaerobic metabolism. Our glucose stimulation results suggested that the balance between the two ATP metabolisms is linked to cell migration, as high cytoplasmic ATP tracks with greater migration. Moreover, our stamp assay results showed another transitional range of oxygen from 4.16-9.14% that promotes the switch from mitochondrial to cytoplasmic ATP metabolism. A dip in mitochondrial ATP coincided with a rise in cytoplasmic ATP levels, at the same range of oxygen levels. However, this hypoxic modulation was absent for high glucose cells, consistent with the results from cell migration in Figure 3a. Similarly, injury-based scratch assay partially recovers this oxygen modulation, where the dip is only seen in the mitochondrial ATP. In the context of a diabetic wound, hyperglycemia may elevate mitochondrial ATP metabolism at the expense of cytoplasmic ATP necessary for cell migration. However, specific levels of tissue hypoxia attempt to recover this cytoplasmic ATP level. This new relationship between hypoxia and glucose suggested a common factor exists between their modulations of cell migration.

Our study revealed an interesting finding regarding the role of ROS in cell migration. Our results indicated that high glucose conditions led to high ROS signal intensity compared to 19 low glucose conditions in both stamp and scratch assays, with and without the average oxygen gradient. The intensity of the signal showed a linear increase from hour 1 to hour 15, followed by a steady state until hour 20, and then a further increase until hour 24 in the absence of the average oxygen gradient. The higher intensity of ROS signal in high glucose conditions compared to low glucose conditions supports the notion that excessive ROS levels can cause oxidative stress, which can be harmful to cells and contribute to diseases. Furthermore, the observed linear increase in ROS signal intensity suggests that the ROS production rate is dependent on time, with a possible relationship between glucose levels and ROS production. The steady-state observed from hour 20 to hour 24 may indicate a balance between ROS production and scavenging, which is crucial for maintaining proper cellular functions. Interestingly, the results also showed that the trend was different for low glucose conditions with normalized intensity near 1 for both assays without the average oxygen gradient and a slightly higher value than the high glucose conditions when the average oxygen gradient was present. These findings suggest that ROS production is dependent on glucose levels and that low glucose conditions may not be sufficient to induce ROS production necessary to inhibit cell migration.

Oligomycin was found to be a mitochondrial ATP synthase inhibitor which triggers the cellular metabolic reprogramming to compensate for the energy deficit. The addition of oligomycin has shown a dip in the mitochondrial ATP levels and an elevation in the cytoplasmic ATP (Figures 8, 9). These results show that HaCaT cells under high glucose conditions while applying oxygen gradient, modified their metabolism to enhance migration in the presence of lower mitochondrial ATP. The role of sodium pyruvate as a scavenger of reactive oxygen species (ROS) involves its ability to directly interact with and neutralize ROS molecules. Sodium pyruvate possesses antioxidant properties, allowing it to counteract the harmful effects of ROS within biological systems (**Ramos-Ibeas et al., 2017**). When ROS accumulate in cells, they cause oxidative stress, leading to cellular damage and dysfunction. Sodium pyruvate acts as a ROS scavenger by reacting with and reducing the level of ROS, such as hydrogen peroxide (H_2_O_2_) (**Wang et al., 2020**). By neutralizing those ROS molecules, sodium pyruvate helps maintain cellular redox balance and prevents oxidative damage to cellular components. One important aspect of sodium pyruvate’s ROS scavenging role is its ability to target ROS production sites, such as mitochondria. As a mitochondrial antioxidant, sodium pyruvate enters the mitochondria and interacts with ROS generated during cellular respiration. This helps protect mitochondrial function and integrity, preventing oxidative stress-related mitochondrial damage. Additionally, sodium pyruvate’s ROS scavenging activity contributes to its broader cytoprotective effects. By reducing ROS levels, sodium pyruvate helps safeguard cellular components, including lipids, proteins, and DNA, from oxidative damage (**Ramos-Ibeas et al., 2017**). This antioxidant action can promote cell viability and overall cell migration. Our study showed that sodium pyruvate lowered the ROS intensity levels especially when we have an actual scratch injury (Figure 10a). The mitochondrial ATP levels were the lowest when we had oligomycin and sodium pyruvate while cytoplasmic ATP was the highest mainly when oxygen gradient was applied (Figure 9). Cytoplasmic ATP was higher with sodium pyruvate and oligomycin than with control (high glucose only) for a stamp (Figures 9a, d), but their effect was observed more when an actual injury was applied (Figures 9b, e). Therefore, oligomycin inhibited the mitochondrial ATP synthase which disrupted the electron transport chain and the process of oxidative phosphorylation. To compensate for the reduced mitochondrial ATP production, cells shifted from aerobic respiration, which uses oxygen, to anaerobic respiration (fermentation), particularly the lactic acid fermentation pathway. HaCaT cell migration was successfully rescued when sodium pyruvate and oligomycin were added to all assays and conditions especially when the oxygen gradient was applied.

Our work provided several significances for diabetic wound healing: Hypoxia is a cue for cell migration and specific levels prompt the transition to enhanced migration. Elevated glucose impairs cell migration but is also dependent on hypoxic cues. Scratch assay in general provided faster but less consistent migration results and differs from stamp assay especially when hypoxia is modulated. These findings have important implications for the treatment of diabetic wounds, which often have poor circulation and reduced oxygen supply. Providing specific oxygen gradients may improve cell migration especially to counteract hyperglycemia. These insights could inform the development of new therapeutics for a variety of diabetic conditions.

## 5. Conclusion

The study discussed the use of microfluidic devices for cell culturing, specifically focusing on HaCaT cells and the impact of varying oxygen concentrations on cell culturing. The study also investigated the effect of hypoxia and diabetes on cell migration using stamp and scratch wound assays. The biochemical pathways responsible for generating mitochondrial and cytoplasmic ATP were also studied and how varying glucose concentrations and an oxygen gradient affect the migration of HaCaT cells. The oxygen gradient microfluidic devices fabricated in our lab provided insights into wound healing in patients with diabetes and other hypoxia-related diseases. The study found that scratch assays had faster migration than stamping, but it was less consistent across the entire wound area. Additionally, high glucose levels in the culture medium negatively impacted the migration of HaCaT cells via changes in metabolites, which showed oxidative stress. Overall, glucose levels significantly impaired cell migration and ATP balance, suggesting a shift from migration to metabolic activities. Finally, the study revealed an interesting finding regarding the role of ROS in cell migration and how high glucose conditions led to an increase in ROS production, which negatively regulated HaCaT cell migration. These findings have important implications for the treatment of diabetic wounds and may inform the development of new therapies for a variety of physiological and pathological conditions. Lastly, a range of hypoxic concentrations between 4.16-9.14% evidently promoted the migration of HaCaT cells, suggesting that hypoxia, as modified by metabolism, is required for proper wound healing. Under high glucose levels with an actual injury, sodium pyruvate and oligomycin elevated the cytoplasmic ATP levels and improved the migration of HaCaT cells especially when gradient oxygen was applied.

## Acknowledgements

This work was funded by the following J.F.L. was supported by National Science Foundation under Grant No 175142, M.O. and K.D. were supported equally by NSF 175142 and the Department of Mechanical Engineering at the University of Michigan Dearborn. M.O. and K.D. were supported by the Department of Mechanical Engineering at the University of Michigan Dearborn.

Any opinions, findings, conclusions, or recommendations expressed in this material are those of the author(s) and do not necessarily reflect the views of the National Science Foundation.

## Data Availability Statement

The raw/processed data required to reproduce the findings of this study are available from the corresponding author upon reasonable request.

